# An objective approach to assess colonic pain in mice using colonometry

**DOI:** 10.1101/2020.12.31.424939

**Authors:** Liya Y Qiao, Jonathan Madar

**Author notes:** Corresponding Author: Liya Qiao, MMRB 5046, 1220 E Broad Street, Richmond, VA 23298, USA.

## Abstract

The present study presents a non-surgical approach to assess colonic mechanical sensitivity in mice using colonometry, a technique in which colonic stretch-reflex contractions are measured by recording intracolonic pressures during saline infusion into the distal colon in a constant rate. Colonometrical recording has been used to assess colonic function in healthy individuals and patients with neurological disorders. Here we found that colonometry can also be implemented in mice, with an optimal saline infusion rate of 1.2 mL/h. Colonometrograms showed intermittent pressure rises that was caused by periodical colonic contractions. In the sceneries of colonic hypersensitivity that was generated post 2,4,6-trinitrobenzene sulfonic acid (TNBS)-induced colonic inflammation, following chemogenetic activation of primary afferent neurons, or immediately after noxious stimulation of the colon by colorectal distension (CRD), the amplitude of intracolonic pressure (A_ICP_) was markedly elevated which was accompanied by a faster pressure rising (ΔP/Δt). Colonic hypersensitivity-associated A_ICP_ elevation was a result of the enhanced strength of colonic stretch-reflex contraction which reflected the heightened activity of the colonic sensory reflex pathways. The increased value of ΔP/Δt in colonic hypersensitivity indicated a lower threshold of colonic mechanical sensation by which colonic stretch-reflex contraction was elicited by a smaller saline infusion volume during a shorter period of infusion time. Chemogenetic inhibition of primary afferent pathway that was governed by Nav1.8-expressing cells attenuated TNBS-induced up-regulations of A_ICP_, ΔP/Δt, and colonic pain behavior in response to CRD. These findings support that colonometrograms can be used for analysis of colonic pain in mice.

## INTRODUCTION

Colonometry in human was first described by Joltrain and colleagues a century ago [1]. In colonometry, the intracolonic pressure (ICP) is recorded by an external manometer or a pressure transducer that is connected to a rectal infusion tube and an infusion system via a three-way connector. A solution (e.g., saline) is infused into the colon continuously and in a constant rate, which results in “stretch-reflex” peristaltic contraction of the colon for periodic attempts to evacuate thereby causing intermittent rises of ICP [2]. Since the 1940s, colonometry has become a routine method in the study of neurologic disturbances in defecation [1,3]. White et al., performed colonometry in patients with various types of neurological lesions and demonstrated that the colonometry gave a picture of the motor activity and sensation of the colon as a whole and reflected the activity of the spinal reflex pathway of the colon [1]. In this clinical study, patients were instructed to report any aberrant sensation such as gas, urge to defecate, or cramp-like pain. The sensation of filling in the colon of normal patients was correlated to lower ICP (20-30 cm of water) and the abdominal pain was reported when higher ICP was recorded (>50 cm of water) [1]. In a group of patients with clinically defined multiple sclerosis (MS), urinary urgency and frequency were their most distressing problems that required hourly urination [3]. Colonometry assessment of these MS patients showed rapid rises of ICP during colonic infusion, resulting in a faster changes in ICP over a certain infusion volume (ΔP/ΔV) [3]. These clinical colonometrical studies suggest that (1) colonic mechanical sensitivity, transduced to stretch-reflex contraction, is reflected by the amplitude of ICP (A_ICP_) and (2) it requires smaller infusion volume, or takes lesser time since the infusion rate is constant (Δt), to increase intracolonic pressures in patients with colonic hypersensitivity such as in MS. In colonometry, colonic reflex contraction produces longer duration of pressure rises while the movement of the diaphragm and abdominal muscle contraction attributed by respiratory excursions causes shorter duration of pressure spikes [2,3].

A decade ago, Larauche et al., performed manometric recordings of rat colon in which ICP was recorded by a pressure sensor implemented inside the colon [4]. The manometric traces of ICP in rats showed that colonic contraction produced longer duration (9.7 s) of pressure rises while abdominal muscle contraction signals resulted in shorter duration (<2 s) of ICP spikes [4]. These ICP patterns corresponded closely to those measured by colonometry in human [2,3]. Traditionally, colonic mechanical pain sensation is assessed by referred abdominal muscle contraction through electromyography (EMG) measurement in response to isobaric colorectal distension (CRD) [5,6]. While EMG in mice is technically challenging due to the small body size, local inflammation caused by implantation of chronic EMG electrodes, and the susceptibility of transgenic mice to highly invasive surgical procedures [7,8]. Measurement of ICP as a result of colon stretch-reflex contraction has advantages of being non-invasive. Subsequent studies by Larauche et al., compared ICP with EMG recordings in mice in assessment of visceral pain [8], however this manometric assessment of colonic mechanical sensitivity by a pressure sensor inside the mouse colon has not been adapted by others. The cost of the miniaturized pressure catheter and the accessibility to the custom-made software scripts used by Larauche et al. could be possible disadvantages.

The beauty of colonometry is its simplicity in measurement of ICP. Guided by clinical studies using colonometry to measure ICP to assess colonic stretch-reflex contractions and the activity of the colonic spinal reflex pathway [1-3], we hypothesize that colonometry is also applicable to experimental mice to measure ICP. In analysis of colonometrograms in conscious mice, we adapted the approaches used in human [1-3] to assess the A_ICP_ and the rapidness of the pressure rising (ΔP/Δt) as measurement of colonic stretch-reflex contractions to indicate colonic mechanical sensitivity. We examined mouse models with chronic colonic hypersensitivity post 2,4,6-trinitrobenzene sulfonic acid (TNBS)-induced colonic inflammation [9-12] or acute colonic pain following noxious CRD [13,14]. We also used transgenic mice due to their popularity in mechanistic studies of colonic sensory disorders. The tetrodotoxin-resistant sodium channel Nav1.8 is expressed by colonic afferent neurons in rats and mice and mediates colonic sensation [15,16] by unmyelinated small-diameter nociceptors and myelinated low-threshold mechanoreceptors (LTMRs) [17]. We used chemogenetic approaches to specifically enhance or reduce the activity of Nav1.8 (haplosufficient)-expressing cells to produce a broad elevation or suppression of the pain sensing pathway including colonic pain. By assessing colonometrograms in these unique mouse models with colonic hypersensitivity, we conclude that colonometry has advantages of being simple, non-invasive, repeatable and objective in determining colonic stretch-reflex contractions as physiological outcomes of colonic mechanical sensitivity in free-moving mice.

## MATERIALS AND METHODS

### Experimental animals

Wildtype and genetically modified C57BL/6 adult male mice (2-3-month old, in-house bred) were used. Animals were housed 2-5 per cage to ensure adequate social environment and numbered by ear clips. Standard husbandry conditions with 12:12-h light cycles and free access to regular food/water were provided. The RC::L-hM3Dq mice (Jax Stock No. 026943) and Nav1.8-Cre mice (a gift of Dr. Sulayman Dib-Hajj, Yale School of Medicine and Dr. John Wood, Wolfson Inst. UK [18]) were crossed to generate Nav1.8;hM3Dq mice for Designer Receptors Exclusively Activated by Designer Drugs (DREADD)-based chemogenetic activation of Nav1.8-expressing cells. The R26-LSL-Gi-DREADD (hM4Di: Jax Stock No. 026219) and Nav1.8-Cre mice were crossed to generate Nav1.8;hM4Di mice for DREADD-based chemogenetic inhibition of Nav1.8-expressing cells. DREADD mice were also mated with wildtype mice to produce non-recombination of DREADD (Wt;hM3Dq or Wt;hM4Di) to serve as genetic background control. The sequences of primers for genotyping Nav1.8-Cre (13Salt and Cre 5a) and wildtype (13Salt and 12A) alleles were: 13Salt: GGAATGGGATGGAGCTTCTTAC; 12A: TTACCCGGTGTGTGCTGTAGAAAG; CRE 5a: CAAATGTTGCTGGATAGTTTTTACTGCC. The sequences of primers for genotyping hM3Dq and hM4Di were from the Jackson Laboratory. Experimental protocols involving animal use were approved by the university Institutional Animal Care and Use Committee (IACUC). Animal care was in accordance with the Association for Assessment and Accreditation of Laboratory Animal Care (AAALAC) guidelines.

### Induction of colonic inflammation

A single dose of 2,4,6-trinitrobenzene sulfonic acid (TNBS: 75 µL of 12.5 µg/µL TNBS in 30 % EtOH) was administered into the mouse colon via a polyethylene (PE)-50 catheter through the anus. The proximal tip of the catheter was 2.5 cm inside from the anus. The mouse tail was lifted for 1 min after TNBS installation to avoid drug leakage from the anus. The same amount of 30 % EtOH as vehicle was used in control animals. TNBS-treated mice developed peak colonic inflammation on day 3-4 and persistent visceral hypersensitivity after inflammation resolution (> 7 days) [9,12,19].

### Colonometry

The apparatus and procedure of colonometry in conscious mice were adapted from those used in human[1-3], and illustrated in Figure 1A. Specifically, a PE-50 tube was inserted inside the mouse colon via anus under light anesthesia (1.5 % isoflurane). The proximal tip of the intracolonic tube was located inside the colon 2.5 cm away from the anus. The intracolonic tube was secured onto the mouse tail. The animal was then placed into a round chamber (approximately 20 cm in diameter) with a gridded bottom to provide dry areas during saline infusion. The distal end of the intracolonic tube was connected to a pressure transducer that was further connected to a syringe on a programmable syringe pump. The pressure transducer was coupled to a bridge amplifier and a computer recording system (AD Instrument, Milford, MA). After the animal was completely awake from anesthesia and was adapted to the environment for at least 30 min, the colon was infused with saline by the syringe pump continuously and in a constant rate (customized as 1.2 mL/h in the present study). The intracolonic pressure traces were acquired and analyzed by LabChart Pro 8 (AD Instrument, Milford, MA). After colonometry, the intracolonic tube was removed. Animal was returned to its home cage.

**Figure 1.**
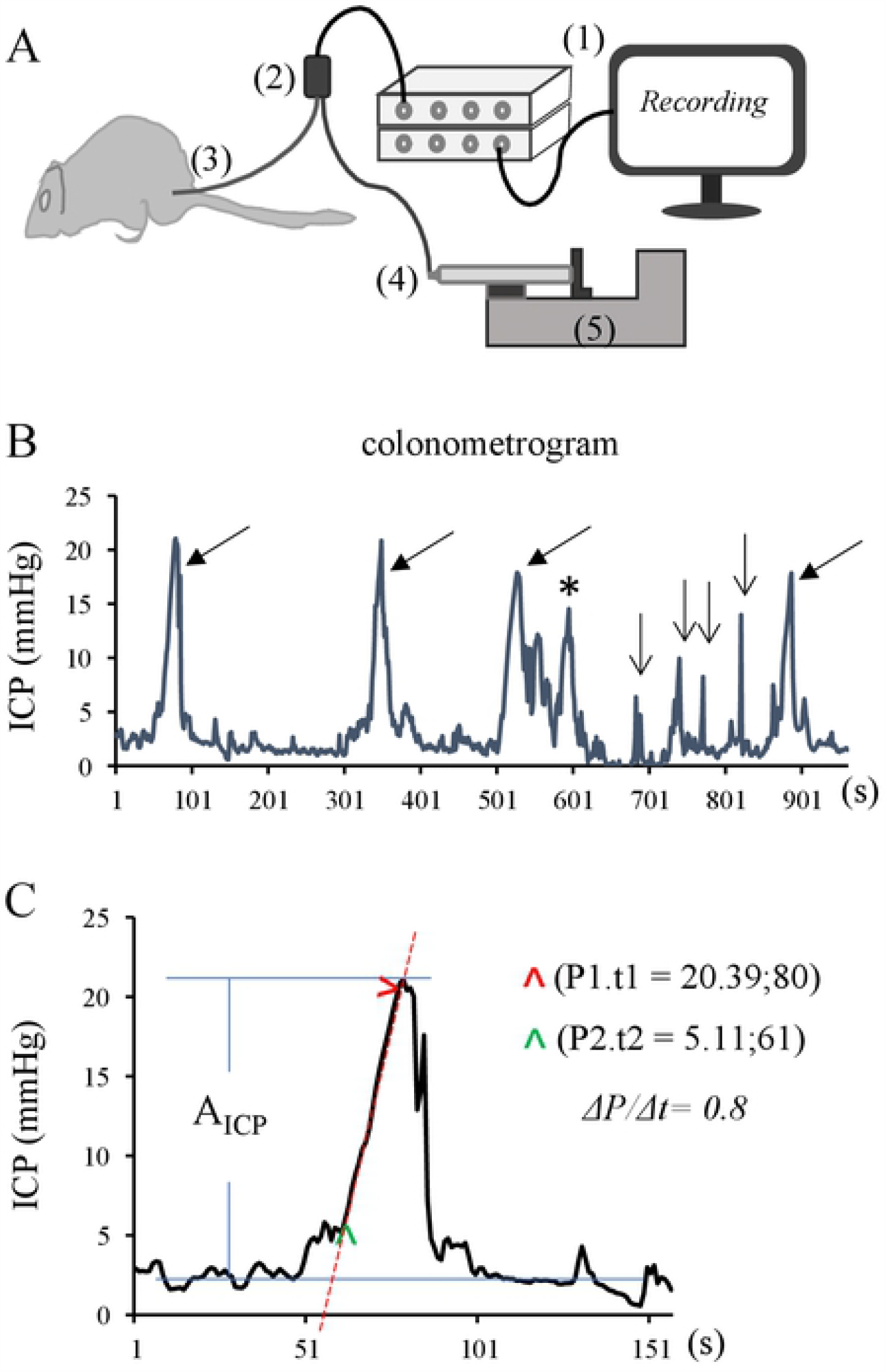
Setup of colonometry in free-moving mice and analysis of colonometrogram. (A): Schematic diagram of the apparatus of colonometry includes (1) a pressure recording system, (2) a pressure transducer, (3) an intracolonic catheter, (4) a syringe, and (5) a programable syringe pump. (B): A representative colonometrogram shows a tracing of intracolonic pressures that include pressure rises as results of colonic stretch-reflex contractions (indicated by arrows) and pressure spikes as results of respiratory excursions (indicated by vee arrows), noting a second wave due to incomplete excretion (indicated by asterisk). (C): Analysis of the amplitude of intracolonic pressures (A_ICP_) and the rapidness of pressure rising (ΔP/Δt).

### Colonometrogram

A representative colonometrogram (Figure 1B) illustrated intermittent pressure rises as results of periodical colonic contractions to attempt to evacuate the saline infused during colonometry. Colonic stretch-reflex contractions-induced intracolonic pressure rises had relatively longer (> 50 s) durations (Figure 1B, indicated by arrows). The pressure spikes with shorter (5-20s) durations (Figure 1B, indicated by vee arrows), deduced from human studies[2], were artifacts caused by respiratory excursions. The pressure rises immediately following colonic contraction (Figure 1B, marked by asterisk) occasionally occurred which might be related to incomplete emptying from the previous colonic contraction. Inferred from human studies with colonometry [1-3] and studies in rodents with manometry of the colon [4,8], we focused on analyzing the pressure rises caused by colonic reflex contractions (Figure 1B, indicated by arrows). The parameters of colonic stretch-reflex contractions that we analyzed, according to the studies in humans [1-3], include (1) determining the amplitude of colonic contraction-induced intracolonic pressure rise (A_ICP_) by subtracting the baseline pressure from the peak pressure (Figure 1C, indicated by blue lines), (2) measuring the distance (time lapse) between adjacent A_ICP_ as inter-contraction intervals (ICI) to reflect, inversely, the frequency of colonic reflex contractions, and (3) calculating the rapidness of pressure rising using linear regression and calculation of the slope of the pressure rising curve (Figure 1C, indicated by a red line) and presented as the ratio of changes in intracolonic pressure (ΔP) over changes in infusion volume (ΔV) or infusion time (Δt) since the infusion rate was constant:

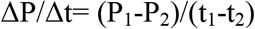

here P represented intracolonic pressure and t represented the time point at which the corresponding intracolonic pressure was recorded (Figure 1C). The area under curve (AUC) was not considered as a readout due to that AUC was positively correlated to the amplitude of the intracolonic pressures and negatively correlated to the rapidness of pressure rising (sharpness of the curve) therefore the physiological meaning of AUC cannot be linked to the strength of stretch-reflex contraction of the colon.

### Immunohistochemistry

Under anesthesia, mice underwent transcardial perfusion by Krebs buffer followed by 4 % paraformaldehyde. The thoracolumbar dorsal root ganglia (DRG) were dehydrated in 25 % sucrose and sectioned at a thickness of 8 µm. The sections were incubated with primary antibodies overnight at room temperature followed by fluorescence-conjugated species-specific secondary antibodies. For rabbit anti-p-CREB (cAMP response element-binding protein) primary antibody (1:2000, Cell Signaling), aminomethylcoumarin (AMCA)-conjugated donkey anti-rabbit secondary antibody (1:100, Jack ImmunoResearch, West Grove, PA) was used. For rabbit anti-p-Akt primary antibody (1:1000, Cell Signaling), Alexa 488-conjugated donkey anti-rabbit secondary antibody (1:500, ThermoFisher Scientific) was used. The stained slides were coverslipped with Citifluor mounting medium and visualized on a ZEISS fluorescent microscope with a multiband filter set. Control sections incubated in the absence of primary or secondary antibody were also processed and evaluated for specificity or background staining levels. The specificity of p-CREB and p-Akt antibodies were also validated by western blot in our previous studies that demonstrated a single immunoreactive band at the correct molecular weight [20,21]. Positively stained cells with visible nucleus were counted. DRG neurons that contained nuclear stain of p-CREB or cytoplasmic stain of p-Akt were analyzed. The number of p-CREB immunoreactive cells was normalized by p-CREB expression in Nav1.8/mCherry-expressing cells. The number of p-Akt immunoreactive cells was normalized by the area of DRG that contained cell bodies (excluding the area of fibers).

### CRD

Under light anesthesia, a mini-balloon coupled to a PE-50 catheter was inserted transanally into the colorectum with the center of the balloon located at a position of approximately 2.5 cm away from the anus. The catheter was secured to the mouse tail. The distal end of the catheter was attached to an empty syringe and a sphygmomanometer pressure gauge via a three-way connector to inflate the mini-balloon and record the intra-balloon pressure. The animal was placed into a clear Plexiglas animal enclosure (IITC) for recovery.

#### Induction of colonic pain by noxious CRD

Immediately after the animal was awake, the intracolonic mini-balloon was rapidly inflated to 80 mmHg and remained for 10 seconds. The mini-balloon was then quickly deflated to 0 mmHg and remained for 5 seconds. This inflation/deflation pattern was repeated 3 times. Sham mini-balloon insertion (0 mmHg intra-balloon pressure) served as control. The paradigm of using noxious CRD to produce acute colonic mechanical pain was adapted from previous publications [13,14] and customized for the present study.

#### Colonic pain threshold responses to CRD

The balloon-catheterized animal was allowed for recovery and exploration of the new environment in the animal enclosure for at least 30 min. The movements/behaviors of the animal were closely observed during a gradual increment of the CRD pressure. The threshold pressure was recorded when the animal demonstrated a sudden immobilization and/or widening of the hind legs, followed by attempts to lick/bite the lower abdomen/anus area (manually interfered the biting to protect the catheter). The tests were performed in a blind manner by which one person was assigned specifically to read the behavior of the animals and gave the signal to the person who read and recorded the pressure gauge. Each animal was tested for 3 times with at least 10-min intervals until the animal restored normal behavior to freely explore the chamber. The values from the 3 trials for each animal were averaged.

### Drug treatment

Suggested by the Jackson Laboratory, the hM3Dq DREADD induces the canonical Gq pathway to depolarize and activate neurons specifically following administration of clozapine-n-oxide (CNO), and the hM4Di DREADD induces the canonical Gi pathway to effectively silence the activity of neurons specifically following administration of CNO. Since 2016, concerns have been raised on the use of relatively high dose of CNO to specifically regulate DREADD. The major concern is its off-target activity by its back-metabolism into clozapine which could occur several hours after CNO injection. Additional studies suggest that intraperitoneal (i.p.) administration of CNO (up to 10 mg/kg body weight) did not cause off-target effects up to 150 min post CNO injection [22]. Thus far, CNO is still the most commonly used chemogenetic actuator in DREADD implementation in vivo. To minimize the off-target effects, we tested the dosage of CNO (Tocris Bioscience) starting from a low dose (1 mg/kg) up and found that 3 mg/kg was adequate in the present study. We also validated the off-target effects of CNO using Wt;DREADD mice and Nav1.8-Cre mice that did not have recombination of DREADD expression. Application of CNO in both DREADDq and DREADDi animals provided further validations. The same amount of DMSO was used as vehicle control for CNO.

### Statistical analysis

At least 3 animals were used for each specific experimental group (see Figure legends for the specific numbers of animals used for each experiment). The raw data from each animal were averaged as one sample (n). GraphPad Prism5 was used for data analysis. The processed data were presented as mean ± SEM. When comparison was made between two groups, Student *t* test was used. For comparison among 3 or more groups, One-way ANOVA with Newman-Keuls Multiple Comparison Test was used. p ≤ 0.05 was considered significant.

## RESULTS

### Effects of infusion rates on colonometrogram

We tested a variety of saline infusion rates for colonometry in wildtype mice. We also observed animal behavioral changes during saline infusion. The infusion rate above 4 mL/h produced pain-like behaviors (hatch-backed, immobilization). At the infusion rate of 2 mL/h, mice exhibited awareness of saline infusion and frequently licked/bit the anus area (a close attention was paid to distract the animals away from biting the tube). At 1.2 mL/h and 0.4 mL/h saline infusion rates, animals did not show any behavioral abnormality. However, at 0.4 mL/h saline infusion rate, the colonic stretch-reflex contractions generated small A_ICP_ (Figure 2A) which was difficult to be extracted from the baseline and from the artifacts caused by respiratory excursions. At 1.2 mL/h saline infusion rate, the A_ICP_ could be clearly extracted out (Figure 2B) thus this saline infusion rate should be optimal for mice. During colonometrical recording, we switched the infusion rates from low to high, and then backwards when testing the optimal infusion rate. We recorded at least four pressure cycles for a designated infusion rate before switching to a different infusion rate. The sequence of application of different infusion rates did not affect animal behaviors as well as the parameters of the colonometrograms such as the values of A_ICP_ and ΔP/Δt for a specific infusion rate. We calculated the values of A_ICP_ (Figure 2A-C) and ΔP/Δt (Figure 2E-G: E was the first contraction from A; F was the first contraction from B; G was the first contraction from C. Dotted lines in E-G were regression lines of pressure rises) up to 2 mL/h and found that the larger infusion rate that caused bigger mechanical forces to the colon produced higher values of A_ICP_ (Figure 2D) and ΔP/Δt (Figure 2H). The values of ICI were not significantly affected when the saline infusion rate was changed (for 0.4 mL/h, ICI=205.3±23.9s; for 1.2 mL/h, ICI=222.7±47.1s; for 2.0 mL/h, ICI=231.3±18.6s; n=4, p=0.85).

**Figure 2.**
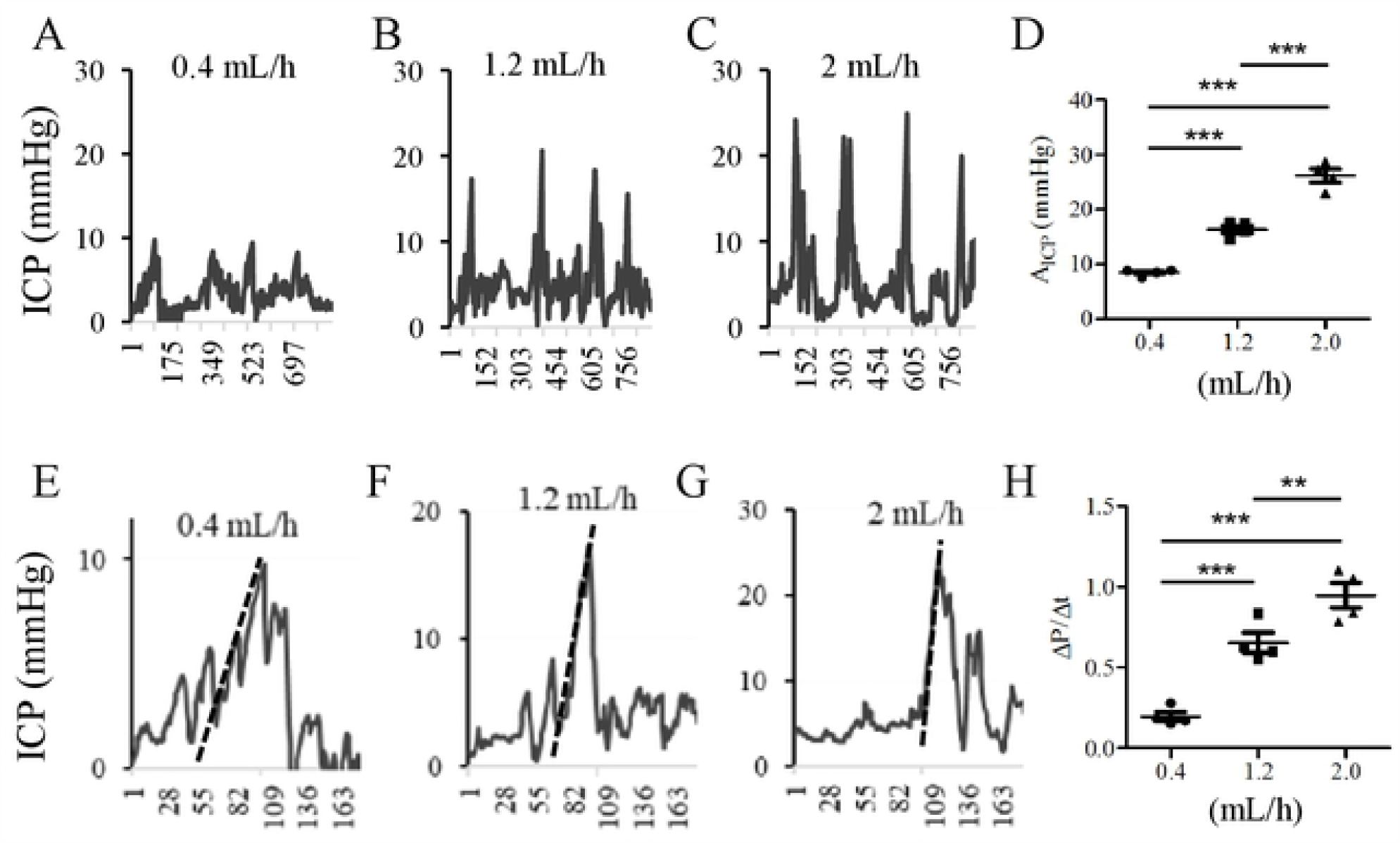
Effects of saline infusion rates on colonometrogram. Colonometrical recordings of wildtype mice under infusion rate of (A): 0.4, (B): 1.2 or (C): 2.0 mL/h. (D): Comparison of A_ICP_ under different infusion rates and the higher rate achieved a larger value of A_ICP_. (E-H): Analysis of the values of ΔP/Δt under the infusion rate of (E): 0.4, (F): 1.2 or (G): 2.0 mL/h and the higher rate achieved a larger value of ΔP/Δt (H). n= 4 animals per condition. ***, p<0.001; **, p<0.01. One-way ANOVA with Newman-Keuls Multiple Comparison Test.

### Colonometrogram in TNBS-induced colonic hypersensitivity

Intracolonic installation of TNBS induces colonic inflammation and postinflammatory colonic hypersensitivity [9-12]. To examine whether colonic stretch-reflex contractions were changed in TNBS-induced colonic hypersensitivity post colonic inflammation, we performed colonometry on day 21 post vehicle (Figure 3A) or TNBS treatment (Figure 3B). We found that TNBS treatment increased both the values of A_ICP_ (Figure 3C) and ΔP/Δt (Figure 3D). We also compared the ICI between vehicle control and TNBS-treated mice and found no significant difference (vehicle: 173.3±49.96s; TNBS: 217±38.45s; n=4, p=0.52). It appeared more occurrence of insufficient emptying in TNBS-treated mice (compare Figure 3B to A, indicated by arrows) in which a second pressure spike immediately followed a previous colonic contraction.

**Figure 3.**
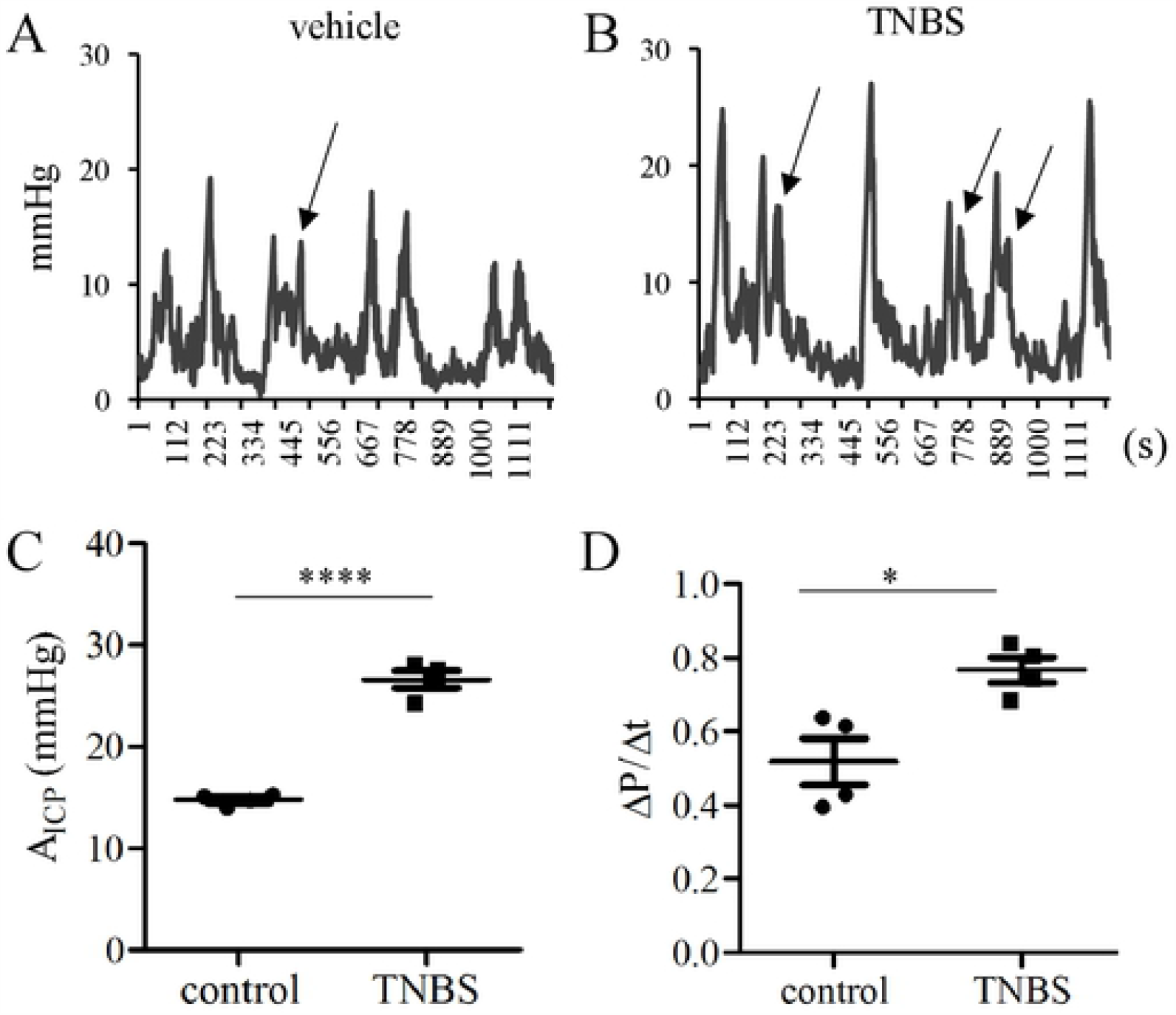
Colonometrical recordings of TNBS-treated wildtype mice. (A): Mice receiving vehicle treatment served as control. (B): Mice were recorded on day 21 post TNBS treatment. (C): Comparison of A_ICP_. (D): Comparison of ΔP/Δt. n= 4 animals per condition. ****, p<0.0001; *, p<0.05. Student *t* test.

### Colonometrogram following Nav1.8-Cre-based chemogenetic activation

Increased expression and channel activity of Nav1.8 in DRG neurons were associated with visceral hypersensitivity as results of colitis, nematode infection, neonatal colonic inflammation, and stress [16]. Here we used chemogenetic approaches to specifically enhance the activity of Nav1.8 - expressing cells to produce a broad elevation of the pain sensing pathway including colonic pain. We exposed Nav1.8;hM3Dq mice to CNO to drive the activation of DRG neurons as well as visceral pain. Nav1.8-Cre-directed expression of Gq-DREADD (hM3Dq) resulted in replacement of green fluorescent protein (GFP) by hM3Dq/mCherry/2ACT88 fusion protein thus having Nav1.8-expressing (haplosufficient) DRG neurons expressed mCherry (Figure 4A and 4B, red cells indicated by arrows; in 4C and 4D, GFP disappeared in these neurons indicated by arrows). CNO treatment of Nav1.8;hM3Dq mice did not change the number of DRG neurons expressing Nav1.8/mCherry (compare Figure 4B to 4A). To examine the effect of CNO treatment on the activity of Nav1.8/mCherry DRG neurons, we measured the expression of p-CREB, a calcium-sensitive molecular switch in neuronal activity [23], in Nav1.8/mCherry DRG neurons 30 min after CNO injection. We found that the immunoreactivity of p-CREB (Figure 4E and 4F, blue nuclear stains, also shown in Figure 4G, 4H as grayscale) in Nav1.8/mCherry-expressing DRG neurons (Figure 4I, merged from 4A, 4C and 4E; 4J, merged from 4B, 4D and 4F; indicated by arrows) was markedly up-regulated following CNO treatment when compared to vehicle (DMSO) treatment (Figure 4K). CNO treatment of Nav1.8-Cre(+/-) mice did not cause p-CREB up-regulation in DRG.

**Figure 4.**
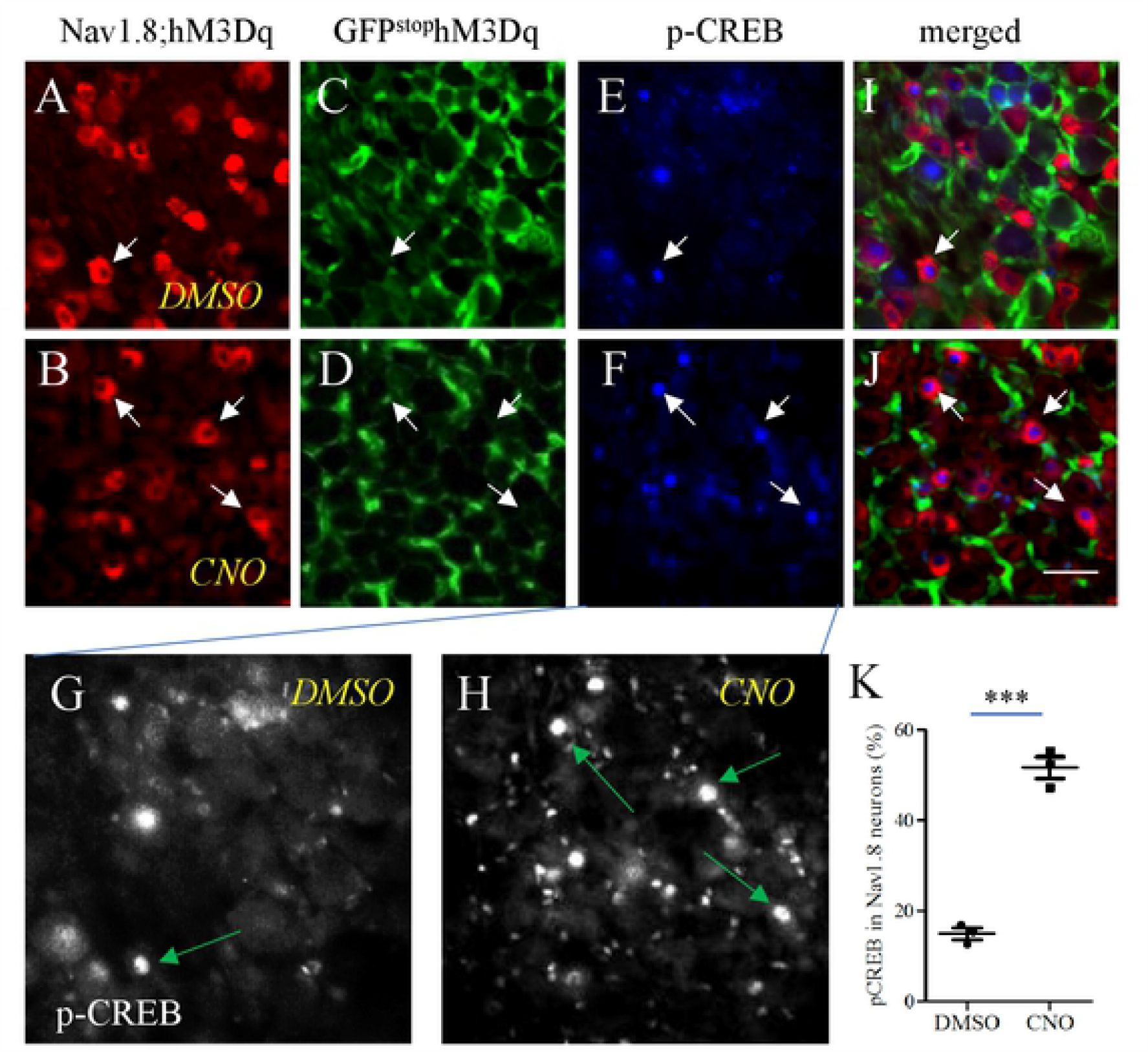
Nav1.8-Cre-driven DREADD-based activation of DRG neurons. Cre-based recombination resulted in mCherry expression (A, B, indicated by arrows) and disappearance of GFP (C, D, indicated by arrows) in Nav1.8-expressing DRG neurons. A subpopulation of Nav1.8-expressing DRG neurons (A, B, red cells) expressed p-CREB (E, F, blue nuclear stain, also shown in Figure 4G, 4H as grayscale, indicted by arrows). The level of expression of p-CREB in Nav1.8-expressing DRG neurons was quantified by their co-localization (I, J, indicated by arrows) and presented as percentage (K). n= 3 animals per condition. ***, p<0.001; Student *t* test. Calibration bar = 50 µm.

We next performed colonometry in Nav1.8;hM3Dq mice 1 hour following vehicle (DMSO) (Figure 5A) or CNO (Figure 5B) injection. CNO treatment significantly increased the values of A_ICP_ (Figure 5C) and ΔP/Δt (Figure 5D) when compared to vehicle control. There was no difference (p=0.84, n=3) in the ICI between DMSO (236.3±22.09 s) and CNO treatment (246.5±45.06 s) of the Nav1.8;hM3Dq mice. We also tested the effects of CNO in Wt;hM3Dq mice. Either the value of A_ICP_ or ΔP/Δt was changed by CNO treatment (A_ICP_: 17.14±0.98 (DMSO) vs 16.95±0.96 (CNO) mmHg, p=0.9; ΔP/Δt: 0.5±0.08 (DMSO) vs 0.52±0.07 (CNO), p=0.9).

**Figure 5.**
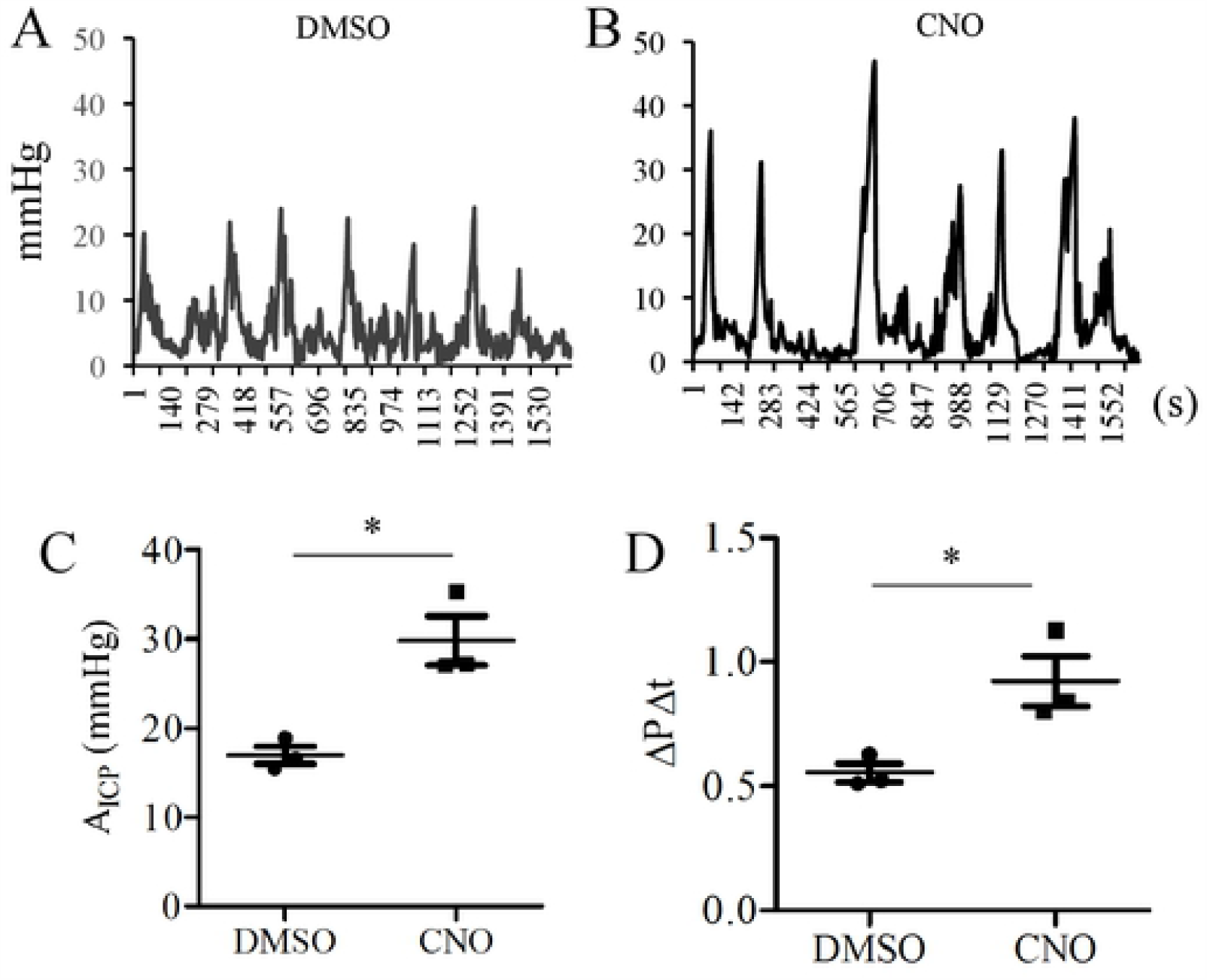
Colonometrical recordings of Nav1.8;hM3Dq mice. (A): Control animals received DMSO treatment. (B): CNO treatment that evokes the activation of Nav1.8-expressing cells alters colonic function. (C): Comparison of A_ICP_. (D): Comparison of ΔP/Δt. n= 3 animals per condition. *, p<0.05. Student *t* test.

### Suppression of TNBS-induced colonic hypersensitivity by Nav1.8-Cre-based chemogenetic inhibition

TNBS-colitis (7-10 days) in mice increased the protein expression and current density of Nav1.8 in DRG neurons post inflammation resolution [15,24]. Previous studies showed that genetic or pharmacological inhibition of Nav1.8 reduced thermal and mechanical hypersensitivity caused by inflammation, nerve injury, or bacterial infection [25-28]. Here we used CNO-induced DREADD-based inhibition of Nav1.8-expressing cells to suppress TNBS-induced colonic hypersensitivity in Nav1.8;hM4Di mice. As shown above, colonic hypersensitivity was associated with the increased values of A_ICP_ or ΔP/Δt following TNBS treatment (Figure 3). We therefore compared the values of A_ICP_ or ΔP/Δt in TNBS-treated Nav1.8;hM4Di mice with or without CNO treatment. On day 21 following TNBS treatment, we performed colonometry 1 hour after vehicle (Figure 6A) or CNO (Figure 6B) injection. The average value of A_ICP_ in TNBS-treated Nav1.8;hM4Di mice was 26.93 ± 0.99 mmHg (Figure 6C), which was similar to the value in TNBS-treated wildtype mice (A_ICP_ = 26.62 ± 0.86 mmHg, referred to Figure 3). CNO treatment decreased the value of A_ICP_ to 18.03 ± 0.91 mmHg in TNBS-treated Nav1.8;hM4Di mice (Figure 6C). The average value of ΔP/Δt was also decreased by CNO in TNBS-treated Nav1.8;hM4Di mice (Figure 6D). There was no change in the ICI following CNO treatment.

**Figure 6.**
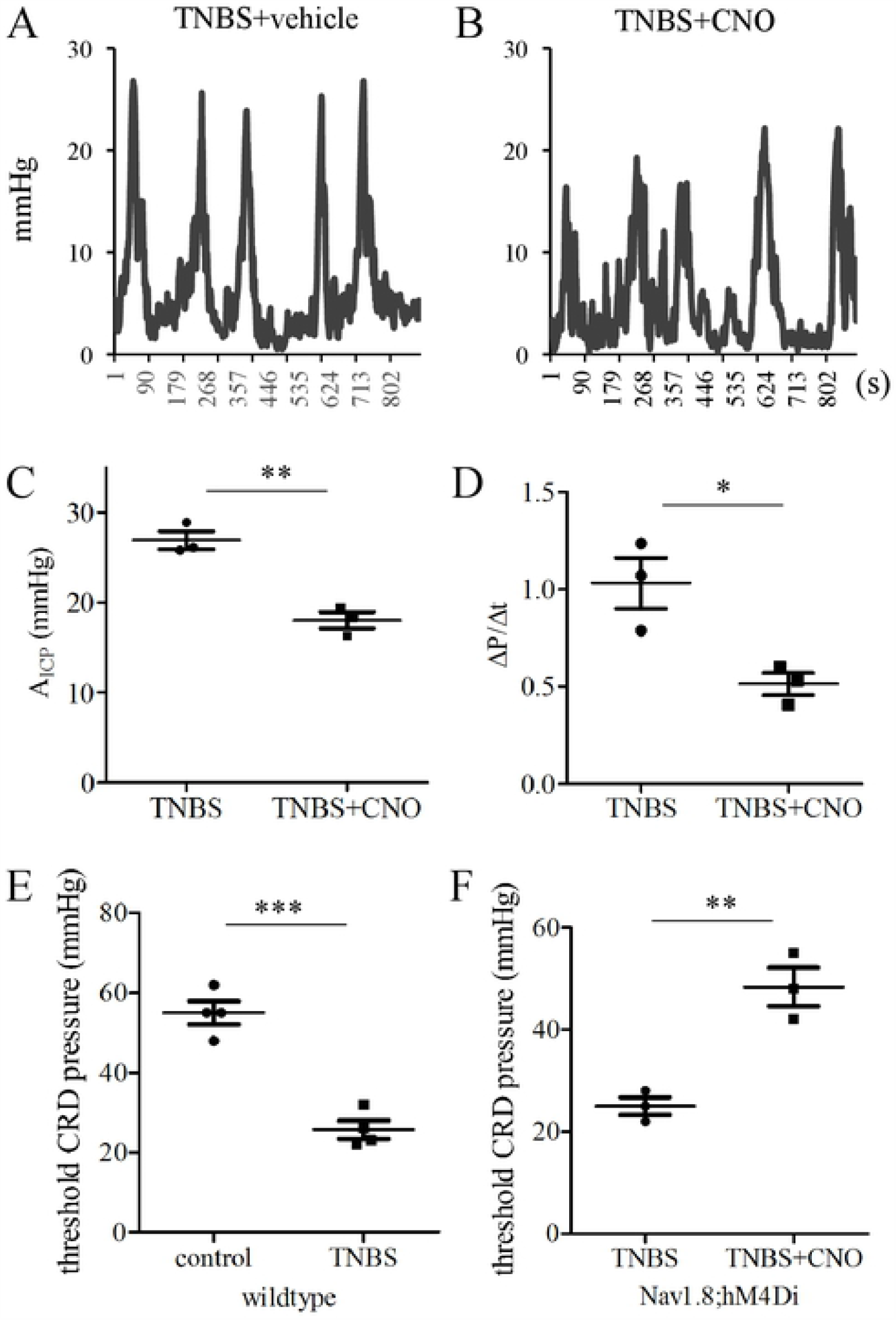
Suppression of TNBS colitis-induced visceral hypersensitivity by chemogenetic inhibition of Nav1.8-expressing cells. Colonometrical recordings of TNBS-treated Nav1.8;hM4Di mice that received DMSO (A) or CNO (B) injection. (C): Comparison of A_ICP_. (D): Comparison of ΔP/Δt. n=3 animals per condition. Colonic pain behavioral responses to CRD in wildtype mice (E: n=4 animals per condition) and Nav1.8;hM4Di mice (F: n=3 animals per condition). ***, p<0.001; **, p<0.01; *, p<0.05. Student *t* test.

For further validation, we performed gross behavioral tests to animals receiving the same treatment. We observed animal behavioral responses to increased gradient of CRD. The threshold pressure for TNBS-treated wildtype mice to demonstrate painful behavior was significantly lower when compared to control (Figure 6E). CNO-induced chemogenetic inhibition decreased the responsiveness of TNBS-treated Nav1.8;hM4Di mice to CRD and increased the pain threshold (Figure 6F).

### Noxious CRD-induced colonic pain

Noxious CRD was used routinely to induce colonic pain in mice that involved an immediate increase in phosphorylated MAPK ERK 1/2 (p-ERK) in the spinal dorsal horn [13]. We found that noxious CRD in wildtype mice also increased the number of thoracolumbar DRG neurons expressing the phosphorylated Akt (p-Akt) within 30 min (Figure 7A-C), suggesting a rapid activation of sensory neurons. In parallel, we found that noxious CRD increased the values of A_ICP_ and ΔP/Δt in colonometrograms of wildtype mice that were examined at 1 h post CRD when compared to sham balloon insertion (Figure 7D-7G). The ICI between sham and CRD was not significantly different.

**Figure 7.**
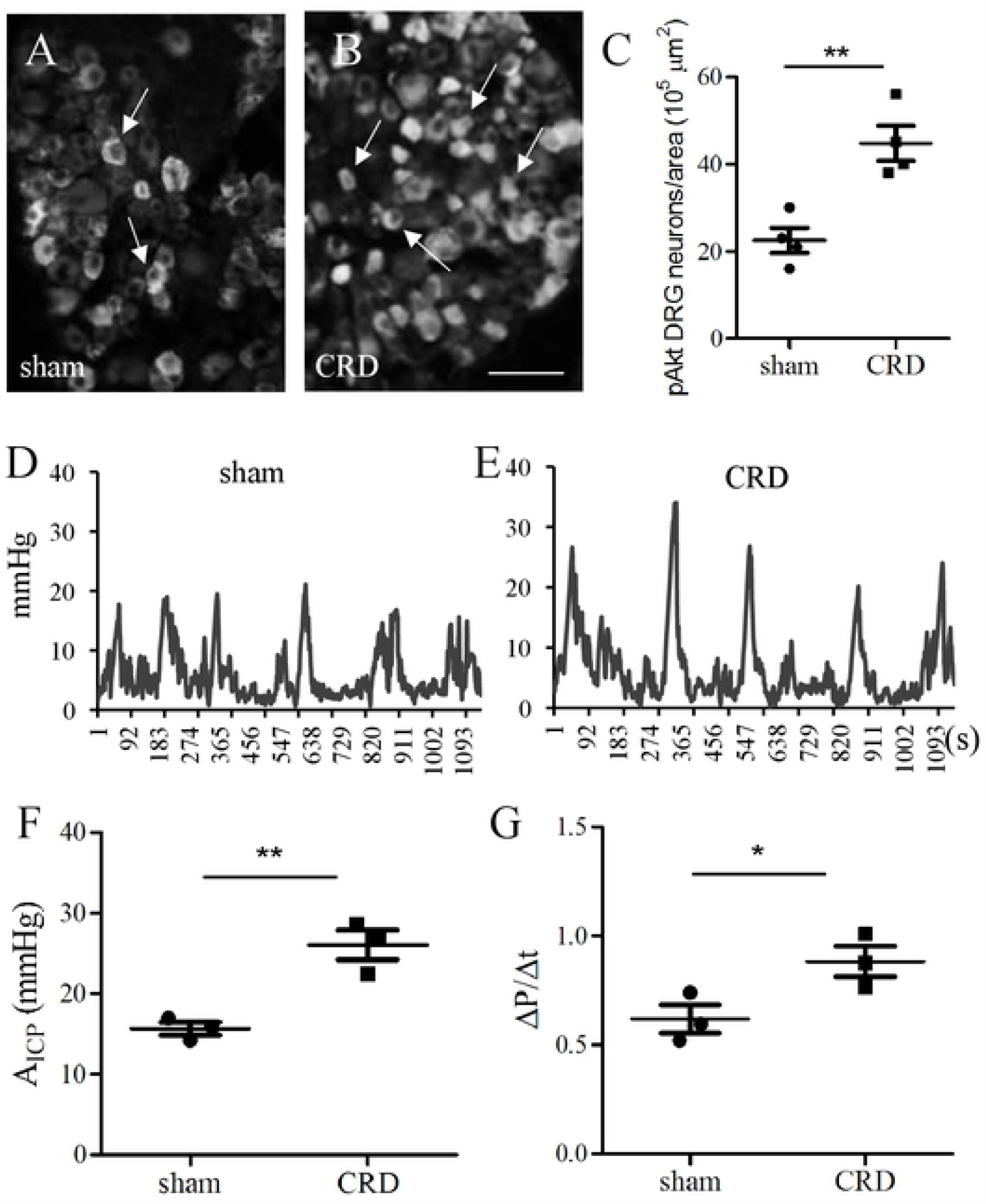
DRG neuron activity and colonometrical recordings following noxious CRD stimulation. (A-C): Immunostaining of p-Akt in thoracolumbar DRG (n=4 animals per condition). (D-E): Colonometrograms from mice that received sham balloon insertion or noxious CRD. (F): Comparison of A_ICP_. (G): Comparison of ΔP/Δt. n=3 animals per condition. **, p<0.01; *, p<0.05. Student *t* test.

## DISCUSSION

Guided by information obtained from clinical settings [1-3], we performed colonometry in mice by infusing saline into the distal colon in a constant rate and simultaneously recorded ICP. Similar as in humans, colonometrograms in mice demonstrated intermittent pressure rises due to stretch-induced peristaltic reflex contractions of the distal colon. We analyzed the colonometrograms for the amplitude of intracolonic pressures (A_ICP_), the rapidness of pressure rising (Δp/Δt), and the colonic inter-contraction intervals (ICI) to determine colonic function in healthy and diseases. We found that the values of A_ICP_ and Δp/Δt were increased in several pathophysiological states that had heightened colonic sensory activity. Examples assessed in the present study included TNBS-induced visceral hypersensitivity, chemogenetic activation of the sensory pathways governed by Nav1.8-expressing cells, and noxious CRD-evoked colonic pain. Furthermore, we found that TNBS-evoked up-regulation in the values of A_ICP_ and Δp/Δt was attenuated by Cre-based chemogenetic inhibition of specific Nav1.8-expressing cells. Inhibition of Nav1.8 also reversed TNBS-evoked colonic pain behaviors in response to CRD. We therefore deduced that the increases in the values of A_ICP_ and Δp/Δt from colonometrograms could be indicators of colonic hypersensitivity. Among all the parameters, the increases in the amplitude of intracolonic pressures are the most straightforward results of increased colonic mechanical sensitivity.

Colonic stretch-reflex contraction is initiated by colonic sensation to the mechanical deformation of the colon wall such as during saline infusion in colonometry. When threshold is reached, the sensory signals received by DRG neurons are conveyed to the spinal reflex pathway by which the motor system is driven to trigger colonic contraction that causes increases in ICP. Excretion of colonic content, e.g., the saline during colonometry, brings down the ICP before starting to build up enough force to elicit another stretch-reflex colonic contraction, forming intermittent pressure cycles. When the colonic afferent neurons are sensitized and hyperactive, such as following TNBS treatment, chemogenetic activation, or noxious CRD stimulation, the sensory responses to colonic stretch (e.g., during saline infusion) are greatly enhanced. The strengthened sensory-motor reflex and increased neurotransmitter release could evoke stronger colonic contraction resulting in higher amplitude of ICP (a larger value of A_ICP_). The surge of ICP caused by colonic reflex contraction could also be rapid due to increased firing thereby resulting in a larger value of Δp/Δt. When the enhanced sensory activity is suppressed such as by chemogenetic inhibition of Nav1.8-expressing cells in TNBS-treated mice, the activity of colonic stretch-reflex pathway and colonic contraction are also reduced therefore the values of A_ICP_ and Δp/Δt are subsequently decreased. In free-moving mice, respiratory excursions could also induce ICP rises due to diaphragm movement, however these changes in ICP are much smaller in its amplitude and duration when compared to stretch-reflex contraction-induced ICP rises thereby they are considered as artifacts in data analysis.

Our results are in line with those obtained from colonometry in humans and colonic manometry in rodents. In human, continuous perfusion of the bowel with saline at a constant flow rate results in waves of ICP at regular intervals [2]. Patients with spinal cord injury demonstrate a rapid rise of ICP recorded by colonometry [1]. The severity of abdominal pain in patients with nerve injury or irritable bowel syndrome is associated with the increased amplitude of ICP recorded [1,29]. MS patients who show bladder hyperactivity and bowel dysfunction also have increased ICP assessed by colonometry [3]. Measurement of ICP as physiological outcomes of colonic sensitivity is also tested in rodents using colonic manometry [4,8]. In manometric assessment of the colon, ICP is sensed by a pressure sensor placed inside the colon to respond to balloon distension. In colonometric assessment of the colon, ICP is acquired by an external pressure transducer connected to an intracolonic tube and is recorded in response to colonic stretch to saline infusion. Although the procedures of these two methods are slightly different, the principles of the technologies are similar in which the colonic stretch-reflex contractions are recorded by ICP to reflect the sensory responses to colonic mechanical stimulations.

To validate colonometry in mice, we choose the well-established TNBS-colitis model for its unique feature of post inflammatory visceral hypersensitivity [12,19]. We find increased values of A_ICP_ and Δp/Δt in TNBS-treated animals. Colonic mechanical pain is mediated by sensory neurons in DRG including nociceptive neurons and LTMRs in which Nav1.8 is highly expressed [17]. When we activate Nav1.8-expressing cells including Nav1.8-expressing DRG neurons by a CNO-induced DREADD-based chemogenetic approach, we find that the values of A_ICP_ and Δp/Δt are also increased. While when we inhibit the activity of Nav1.8-expressing cells, TNBS colitis-evoked up-regulation of A_ICP_ and Δp/Δt is attenuated, meanwhile TNBS colitis-induced pain behavior in response to CRD is also suppressed. These results suggest a correlation of the values of A_ICP_ and Δp/Δt to colonic pain. In addition to DRG neurons [17], Nav1.8 expression is also identified in the colon, brain, and heart [16,30]. Thus, chemogenetic activation or inhibition of Nav1.8-expressing cells does not preclude the effects of CNO in the colon and brain to affect colonic mechanical sensitivity. In addition, we find that the values of A_ICP_ and Δp/Δt are also increased following noxious CRD stimulation of the colon, by which the sensory reflex pathways are activated [13]. Collectively, we find consistent results of colonometrograms from mice with colonic hypersensitivity that is induced by different methods.

The approaches used in clinical settings to assess colonic pain and pain in general involve questionnaires and numerical scaling that are not applicable to experimental animals [31,32]. Over the years, several methods have been developed and some of them have been widely used in rats to assess colonic pain. These methods include assessment of abdominal withdrawal reflex (AWR) in response to CRD [10,33-35] and measurement of visceromotor responses (VMR) by EMG [36,37]. However, these methods are not very reliable in mice due to the nature that mice are more active for AWR analysis, and the transgenic mice are more susceptible to surgeries. Recent years a concept of measurement of ICP as physiological outcomes of colonic sensation in mice has been developed. In addition to colonic manometry that was created by Larauche et al. [8], O’Mahony proposed to measure the pressure spikes above the CRD pressure detected by the intracolonic mini-balloon as colonic contractile pressure [34]. However, these techniques have not been widely applied to the research field of visceral hypersensitivity. Adapted from human studies to measure ICP by colonometry [1-3], we for the first time present the feasibility of colonometry in mice to measure ICP as an outcome of colonic stretch-reflex contraction to assess colonic mechanical sensitivity. Infusion of a constant rate of saline into a hallow organ while measuring intra-organ pressures to assess the organ function has been used in pulmonary [38] and urodynamic [39,40] studies, and here in study of colonic function.

## Grant Support

NIH R01 DK118137; CHRB 236-06-18; CTSA 5KL2TR002648

## Disclosures

The authors declare no conflict of interest.

